# High microsatellite but no mitochondrial DNA variation in an invasive Japanese mainland population of the parasitoid wasp *Melittobia sosui*

**DOI:** 10.1101/2024.07.19.604378

**Authors:** Jun Abe, Jun-ichi Takahashi, Koji Tsuchida

**Author notes:** Correspondence: Jun Abe, Faculty of Sciences, Kanagawa University, Kanagawa 221-8686, Japan.

## Abstract

Theory predicts that bottleneck events reduce genetic diversity in invasive populations. The parasitoid wasp *Melittobia sosui* was only identified in the subtropical area of Japanese south islands and Taiwan, but recently found also in the temperate area of the Japanese mainland. The species may expand their distribution northward due to factors such as recent global warming. We investigated population genetics in both the native and invasive areas using mitochondrial and nuclear microsatellite DNA. As expected, there was mitochondrial variation in the native area, but not in the invasive area, which only had one haplotype. However, the two areas had a similar level of microsatellite variation, with averagely 43% and 38% alleles uniquely found in the native and invasive populations, respectively. The difference of genetic variation between mitochondrial and microsatellite DNA in the invasive populations may be explained by the faster mutation rate of microsatellites, as well as the population structure of *Melittobia*, in which subdivided small inbreeding lineages may facilitate the accumulation of mutations. The high proportion of private alleles suggests that the mainland population diverged from the native populations at least 100 years ago, ruling out the possibility that the mainland population was established recently. Instead, the present study suggests that *M. sosui* might have already existed in the mainland but with a low frequency, or the mainland population was derived from a third population which diverged from the native populations over 100 years ago.

## Introduction

How populations are established in novel habitat areas outside the distribution range of the species is one of the fundamental questions in evolutionary ecology and conservation biology. The distributions of species are changing at accelerating rates due to natural processes as well as human activities (Pecl et al. 2017). Recent climate changes such as global warming have leaded to expand the distribution ranges mainly toward higher latitudes in a wide range of organisms (Deutsch et al. 2008; Walther et al. 2009). Human transportations have facilitated organisms to expand regions beyond their natural migration ranges (Hulme 2009; Banks et al. 2015).

Through invasion processes, genetic diversities in invasive populations could become differentiated from ones in native populations. Due to bottleneck events, with an only fraction of the genetic variants transferred from the native habitats by founders, invasive populations often show lower genetic diversity relative to native populations (Nei et al. 1975). However, multiple introductions and introductions by a large number of founders can prevent the reduction of genetic diversity, resulting in even maintaining genetic diversity to the level comparative to the native populations (Dlugosch and Parker 2008; Uller and Leimu 2011). Along with variation introduced by founders, variants can arise by mutations after divergence (Talla et al. 2019; Kaňuch et al. 2021). Although it takes a reasonable period for mutations to occur and contribute to the genetic diversity in the population, the effect of mutation is likely to be more pronounced in genomic regions where mutation rates are faster. In addition, infection by endosymbiotic bacteria *Wolbachia*, which infect a wide range of arthropods and nematodes, has frequently been reported to affect the genetic structure and diversity of their host species (Werren et al. 2008). *Wolbachia* are maternally transmitted, and often manipulate the reproduction of their hosts by the ways such as parthenogenesis, feminization, male-killing, and cytoplasmic incompatibility (Hurst 1993; Werren 1997; Stouthammer et al. 1999). Such reproductive manipulation enhances *Wolbachia* lineages, coupled with maternally co-inherited associated mitochondrial haplotypes, to be vertically transmitted and spread in host populations. Therefore, *Wolbachia*-induced sweep causes reduction in mitochondrial variation but has little effect on nuclear variation. Studies have documented reduced mitochondrial variation in *Wolbachia*-infected populations in variable insect species (Hale and Hoffmann 1990; Werren et al. 2008; Raychoudhury et al. 2010; Miyata et al. 2017).

*Melittobia sosui* is a parasitoid wasp that attacks the larvae and pupae, mainly prepupae, of a variable range of solitary wasp and bee species nesting above ground (Matthews et al. 2009). The distribution of *M. sosui* has been confirmed only from Japanese south islands (Okinawa and Iriomote islands) and Taiwan (Fig. 1) in the subtropical area, but not in the Japanese mainland in the temperate area (van den Assem and Maeta 1980; van den Assem et al. 1982; Maeta 1985; Maeta et al. 1996). However, it has repeatedly been collected from Shizuoka and Kanagawa in the Japanese mainland after 2008 (Fig. 1; J. A. unpublished data). The reproduction of *Melittobia* species is characterized by repeated inbreeding, as mating occurs between individuals that developed on the same host before female dispersal, and related females from the same host often lay eggs together on other hosts within the same host patches (Matthews et al. 2009; Abe and Pannebakker 2017; Abe et al. 2021). Only when occasionally females dispersed from different host patches lay eggs on the same host, outcrossing occurs in their offspring generation (Abe et al. 2021). *Melittobia* is haplodiploid like other Hymenopteran species. Whereas deleterious mutations are hidden in diploid females, they are purged by selection in haploid males, leading to less reduction in fitness by inbreeding depression (Werren 1993). Moreover, unlike many other Hymenopteran species, as chalcid wasps including *Melittobia* have a sex determination system other than the complimentary sex determination (CSD), they do not incur genetic load by producing sterile diploid males as a consequence of inbreeding (Beukeboom and Perrin 2014). Therefore, even if only a single mated female invades in a new habitat area, and conditions in the environment are suitable, she is likely to establish a new population without suffering from inbreeding depression. Because the host species build their nests in the gaps and crevices of not only natural structures but also artificial ones, hosts parasitized by *M. sosui* may be immigrated by human transports. Additionally, there is a jet stream blowing from southwest to northwest in the area. Recent global warming has been reported to cause the northward expansion of various insect species from southern regions in the Japanese archipelago (Kiritani 2006). Therefore, the most straightforward and plausible scenario could be that the newly found *M. sosui* population in the Japanese mainland has recently been established by an immigrant(s) from the southwest area.

**Fig. 1.**
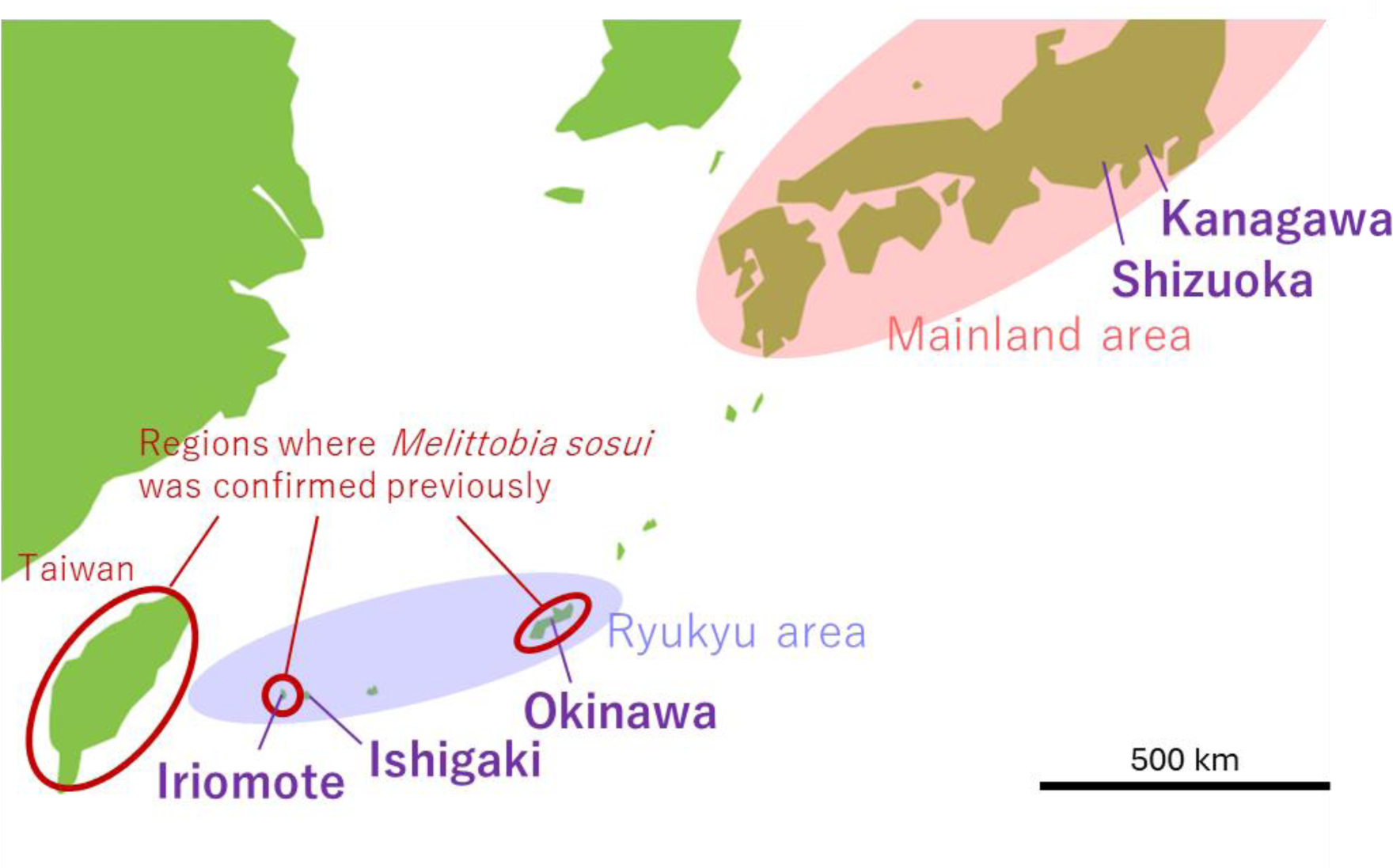
Map around Japanese mainland and Ryukyu areas and Taiwan. *Melittobia sosui* was previously reported from Okinawa, Iriomote, and Taiwan. The present study sampled *M. sosui* from the native area, Iriomote, Ishigaki, and Okinawa in Ryukyu area as well as the new habitat, Shizuoka and Kanagawa in mainland area.

In this study, we examined the genetic structure and variation of *M. sosui* at the levels of nuclear and mitochondrial DNA in both the native area (Ryukyu area) and the new habitat area (Mainland area; Fig. 1). We expected reduced genetic diversity in the new habitats in both nuclear and mitochondrial DNA, if gene flow has been taken place recently from the native area. For the nuclear DNA marker, most of microsatellite markers developed for another *Melittobia* species did not apply for the present species, we newly developed microsatellite markers for *M. sosui* with the method using the next-generation sequence (Abe and Pannebakker 2017). Moreover, as *Wolbachia* might affect the genetic structure and diversity of the populations examined, we also investigated the infection of *Wolbachia* in the populations of *M. sosui*. Finally, based on the results, we discuss the process of how *M. sosui* was introduced and settled in the Japanese mainland and consequent changes in genetic diversity during subsequent population expansion.

## Materials and Methods

### Sample collection and DNA extractions

We conducted field works in the native Ryukyu area comprising Iriomote, Ishigaki, and Okinawa populations and the invasive Mainland area comprising Shizuoka and Kanagawa populations (Fig. 1). We set bamboo traps in Iriomote island for 2008 – 2015, in Ishigaki island for 2008 – 2015 and 2021 – 2023, in Okinawa island for 2008 – 2009, in Fukuroi city, Shizuoka for 2008, and in Hiratsuka city, Kanagawa for 2011 – 2023 (Abe and Pannebakker 2017; Abe et al. 2021). The bamboo canes nested by solitary wasps and bees were brought back to the laboratory, where parasitism by *Melittobia* species was inspected. The juveniles of *Melittobia* were incubated in 25 °C to allow to emerge. After the species were identified, the emerged individuals were preserved in 99.5% ethanol.

Total genomic DNA was extracted using a DNeasy Blood & Tissue Kit (Qiagen) according to the manufacturer’s protocol. For the polymorphism check of the microsatellite markers developed, we examined 6, 10, 7 individuals from Iriomote, Ishigaki, and Kanagawa populations, respectively (Table S1). For the mitochondrial and microsatellite analyses of population genetics, we added additional samples, and totally examined 14, 25, 3, 1, and 10 individuals from Iriomote, Ishigaki, Okinawa, Shizuoka, and Kanagawa populations, respectively (Table S1). We sampled only one individual per bamboo trap to avoid collecting related individuals (Abe et al. 2021). All the individuals were female except one from Okinawa population, as only several male individuals emerged from the brood. As *M. sosui* is haplodiploid and males are haploid, we calculated population genetic estimates that are based on heterozygote frequencies, such as *F*_IS_ and *F*_ST_, with only female individuals discarding the male individual.

### Development of microsatellite DNA markers

DNA was extracted from 20 *M. sosui* individuals of a laboratory strain originated from individuals collected Hiratsuka, Kanagawa in 2017. Whole bodies were digested in 500 μl of 10 mM Tris–HCl (pH 8.0) supplemented with 10 mM EDTA, 0.5% SDS, and 0.5 mg/ml proteinase K at 55 °C for 24 h. Genomic DNA was extracted from the solution once with phenol, once with phenol/chloroform (1:1), and once with chloroform/isoamyl alcohol (24:1). It was then precipitated with ethanol/3 M sodium acetate (20:1), washed with 70% ethanol, dried, and dissolved in TE buffer. DNA samples were stored at −20 °C until further use.

Microsatellite DNA motifs were sequenced with Illumina’s MiSeq platform (Illmina). The quantification of DNA solutions was conducted utilizing the Synergy LX (BioTek) and the QuantiFluor dsDNA System (Promega). The fragmentation of DNA was conducted using a Covaris device under conditions that resulted in the generation of DNA fragments with a length of 500 base pairs. Library preparation was conducted in accordance with the manufacturer’s instructions, utilizing sheared DNA (50 ng) and the KAPA Hyper Prep Kit. The libraries were subjected to 8 cycles of polymerase chain reaction (PCR) amplification. Adapters from the FastGene Adapter Kit (FastGene) were employed. The quantification of the libraries was conducted using the Synergy H1 and the QuantiFluor dsDNA System to ascertain the concentration of the prepared libraries. The quality of the libraries was evaluated using the Bioanalyzer and the High Sensitivity DNA Kit (Agilent Technologies). Sequencing analysis was performed on the MiSeq under 2 x 300 bp conditions using the prepared libraries. The raw sequence data were stripped of adaptor sequences and low-quality bases using default parameters. The data was adjusted through quality filtering using Sickle. Bases with a quality value below 20 were removed, and reads along with their paired reads were discarded if they fell below 120 bases. Clean reads were obtained by assembling high-quality reads using Spades ver 3.10.1.

The reads containing microsatellite regions from the raw sequence data were screened using MSATCOMMANDER program (Faircloth 2008). Selection criteria of microsatellite motifs were at least 15 consecutive complete dinucleotide repeat sequences or at least 10 consecutive complete trinucleotide repeat sequences. Microsatellite primers were designed to be at least 20 bases away from the microsatellite region and at least 100 bp when amplified by PCR. Microsatellite primers were designed using the Primer3Plus software (Untergasser et al. 2007) with these settings. To check amplification and polymorphism for each primer pair designed, PCR was conducted using the M13-tails technique (Schuelk 2000), in a total volume of 5 μl, containing 2.5 μl 2× Type-it Multiplex PCR Master Mix (QIAGEN), 0.5 μl M13 primer (10 μM) labeled a fluorescent dye (6-FAM, VIC, NED, or PET), 0.5 μl 10× primer mix (0.05 μM forward primer with M13 tail and 0.2 μM reverse primer), and 1.5 μl genomic DNA, with the temperature profile of 5 min at 94 °C, then 30 cycles of 30 s at 94 °C, 45 s at 60 °C and 45 s at 72 °C, followed by 8 cycles of 30 s at 94 °C, 45 s at 53 °C and 45 s at 72 °C, and a final extension of 10 min at 72 °C. The PCR products were electrophoresed with an ABI 3130 sequencer (Thermo Fisher) and analyzed with the Peak Scanner software version 1.0 (Thermo Fisher).

Tests for linkage disequilibrium between each locus pair were calculated using the software GENEPOP version 4.7 (Rousset 2008), and significance thresholds were corrected using the false discovery rate (FDR; Benjamini and Hochberg 1995). As a large genetic differentiation was observed between the Ryukyu and Mainland population groups (as shown below), this analysis was separately done for each population group.

### Mitochondrial DNA analysis

PCR was performed to amplify the mitochondrial cytochrome oxidase subunit I (COI) gene region using the primer pair LCO1490 (5′-GGTCAACAAATCATAAAGATATTGG-3′) and HCO2198 (5′-TAAACTTCAGGGTGACCAAAAAATCA-3′) (Folmer et al., 1994). The PCR reactions were conducted by an ABI 2720 thermal cycler (Thermo Fisher), in a total volume of 10 μl, containing 1 μl Ex Taq buffer, 0.8 μl dNTP mixture (2.5 μM each), 0.1 μl Ex Taq Hot Start Version (TaKaRa Bio), 2.5 μl each primer (2 μM), and 1 μl genomic DNA, with the temperature profile of 5 min at 96 °C, then 30 cycles of 30 s at 94 °C, 45 s at 50 °C and 45 s at 68 °C, and a final extension of 2 min at 72 °C. After the PCR products were treated with ExoSAP-IT Express (Thermo Fisher), sequencing reaction was conducted for both strands with the same primers as the PCR using a BigDye Terminator v3.1 Cycle Sequencing Kit (Thermo Fisher). The sequencing products were purified with Gel Filtration Cartridge (Edge BioSystems), and the products were analyzed using an ABI 3130 sequencer (Thermo Fisher).

Assembles for both strands were performed using GeneStudio Professional, version 2.2.0.0 (GeneStudio Inc.), and multiple sequence alignments were done using the ClustalW program (Thompson et al. 2003) as implemented in MEGA11 version 11.0.13 (Tamura et al. 2021). Sequences for three nucleotides which could not be determined unambiguously were excluded, and remaining 532 bp were used for the analyses below. With the sequence data, a haplotype network was constructed using the TCS network implemented in PopART version 1.7 (Leigh and Bryant 2015). To investigate the level of mitochondrial diversity, number of haplotypes, haplotype diversity, and nucleotide diversity were calculated using DnaSP version 6.12.03 (Rozas et al. 2017).

### Microsatellite DNA analysis

PCR was conducted with the same condition as amplification and polymorphism checks above using primer pairs developed for microsatellite loci. The observed and expected heterozygosities (*H*_O_ and *H*_E_), the inbreeding coefficient (*F*_IS_) according to Weir and Cockerham (1984), and the deficiency of Hardy-Weinberg equilibrium (HWE) were calculated using GENEPOP version 4.7 (Rousset 2008) and Arlequin version 3.5.2.2 (Excoffier and Lischer 2010). Pairwise *F*_ST_ values between populations were estimated using Arlequin. To investigate genetic differentiation between the native Ryukyu group and the invasive Mainland group, analysis of molecular variation (AMOVA) was carried out using Arlequin.

To estimate the population genetic structure across all the area, Bayesian clustering analysis was performed using STRUCTURE version 2.3.4 (Pritchard et al. 2000). For the admixture model, the predefined number of genetic clusters to assign individuals (*K*) was set from 1 to 6, and ten independent runs were repeated for each value of *K*. Each run involved 1,000,000 Markov chain Monte Carlo iterations after a burn-in period of 100,000 iterations. The most likely value of *K* was selected based on the method of D*K* (Evanno et al. 2005) using the STRUCTURE HARVESTER (Earl and vonHoldt 2012). The runs of the most likely *K* were summarized using CLUMPAK (Kopelman et al. 2015). To investigate the population structure over the area, phylogenetic trees were also constructed with the neighbor-joining (NJ) method (Saitou and Nei 1987) using Nei’s *D*_A_ genetic distance in POPTREE2 (Takezaki et al. 2010), and graphically edited using MEGA11.

The genetic diversity indices, that is number of alleles, gene diversity, and allelic richness, were calculated for each population and for the Ryukyu and the Mainland population groups using FSTAT version 2.9.4 (Goudet 1995). The number and frequency of shared and private alleles between Ryukyu and Mainland groups were manually calculated from the number of alleles for each locus.

### PCR detection for Wolbachia infection

To investigate the infection of *Wolbachia*, diagnostic PCR was performed to amplify *Wolbachia* surface protein (*wsp*) gene region using the universal primer set of wspF (5′-GGGTCCAATAAGTGATGAAGAAAC-3′) and wspR (5′-TTAAAACGCTACTCCAGCTTCTGC-3′) (Kondo et al. 2002; Abe et al. 2003). The composition of the reaction mix was the same as one for mitochondrial COI above, and the temperature profile consisted of 2 min at 94 °C, then 30 cycles of 1 min at 94 °C, 1 min at 50 °C and 1 min at 70 °C. As positive control, the same PCR was conducted with an individual from a laboratory strain of *Melittobia australica*, which was confirmed to be infected by *Wolbachia*. The PCR products were electrophoresed with agarose gels, and the amplification of about 0.6 kb segment was checked.

## Results

### Development of microsatellite DNA markers

A total of 1,341,634 raw reads were obtained from a *M. sosui* gDNA library by Illumina paired-end sequencing. Microsatellite markers were successfully designed for 32,315 and 1,927 primer pairs that amplify the microsatellite regions of dinucleotide and trinucleotide repeat, respectively. We selected 112 primer pairs of trinucleotide repeat to avoid overlap of alleles due to stutter artifact in dinucleotide repeat. Out of those, 101 loci were successfully amplified, and 65 loci had polymorphism. Of these loci with polymorphism, we selected 55 loci analysis below (Table S2). Tests for linkage disequilibrium showed that there was no linkage disequilibrium in Ryukyu group, while two pairs of loci (Mso007–Mso024 and Mso024–Mso045) were significantly associated with each other (Table S3). As the P value of the other pair of the loci (Mso007–Mso045) was also small and fairly close to the corrected significance threshold, we discarded the two loci Mso024 and Mso045 and used remaining 53 loci for further analyses below (Table S2).

### Mitochondrial DNA analysis

In total, we found 9 unique mitochondrial haplotypes containing 18 polymorphic sites. The haplotypes were split into two discrete clades with at least11 nucleotide differences (Fig. 2). The individuals of the populations from Ryukyu group were distributed into the two clades, while all individuals from Mainland populations shared the same haplotype. The haplotype of Mainland individuals was the most frequent one (Hap1 in Fig. 2) sharing with individuals from Ryukyu populations. Hence, haplotype diversity and nucleotide diversity in Ryukyu group were positive, indicating that there was variation in mitochondrial DNA, but Mainland group had no variation (Table 1). These results correspond to the hypothesis that Mainland populations were originated from a part of individuals from Ryukyu populations.

**Fig. 2.**
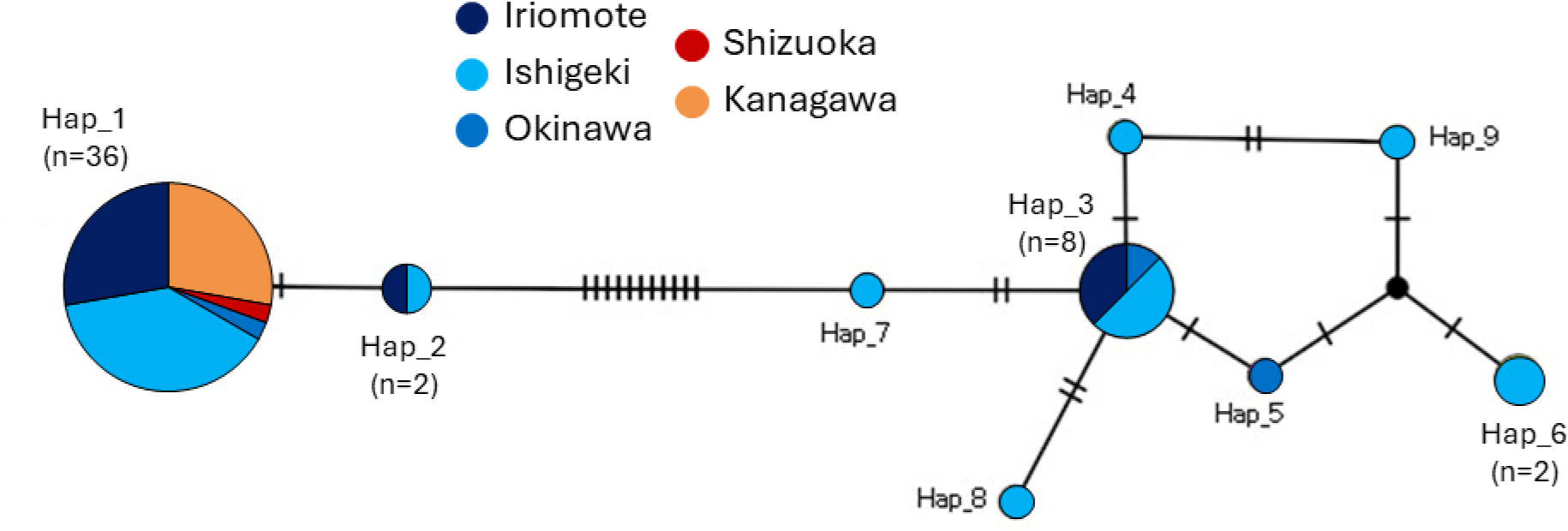
TSC haplotype network of *Melittobia sosui* based on mitochondrial COI sequences. Size of circles is proportional to the number of individuals sampled: dark blue, Iriomote; light blue, Ishigaki; blue, Okinawa; red, Shizuoka; orange, Kanagawa. The number of crossbars on blanches represents the number of mutational changes between haplotypes.

**Table 1.** Genetic diversity indexes of mitochondrial COI haplotypes in *Melittobia sosui*.

| Population<sup>1</sup> | Number of<br>individuals analyzed | Number of<br>haplotypes | Haplotype (gene)<br>diversity | Nucleotide diversity |
| --- | --- | --- | --- | --- |
| per population |  |  |  |  |
| Iriomote | 14 | 3 | 0.473 | 0.0097 |
| Ishigaki | 25 | 8 | 0.673 | 0.0137 |
| Kanagawa | 10 | 1 | 0 | 0 |
| per group<sup>2</sup> |  |  |  |  |
| Ryukyu | 42 | 9 | 0.617 | 0.0127 |
| Mainland | 11 | 1 | 0 | 0 |

<sup>2</sup>Ryukyu group consists of Iriomote, Ishigaki, and Okinawa populations, and Mainland group consist of Shizuoka and Kanagawa populations.

### Microsatellite DNA analysis

The observed heterozygosity *H*_O_, expected heterozygosity *H*_E_, and inbreeding coefficient *F*_IS_ were calculated for loci with polymorphism and for populations with sufficient sample sizes. Overall, inbreeding coefficient *F*_IS_ was high, and 6 out of 12, 10 out of 12, and 5 out of 15 loci in Iriomote, Ishigaki, and Kanagawa populations, respectively, showed significant departure from HWE (Table S4). The *F*_IS_ value calculated over all the loci for the populations was 0.816, 0.798, and 0.557, respectively, and significant departure from HWE was detected for all the populations (all *P* < 0.001). This result is reasonable, given the life history of *Melittobia* with repeated inbreeding (Matthews et al. 2009; Abe and Pannebakker 2017; Abe et al. 2021). All the estimates of pairwise *F*_ST_ between the populations were also high, and significantly deviated from zero, although the pairwise *F*_ST_ values were relatively higher between the native and invasive populations than within native populations (Table. S5). The results of AMOVA, in which Ryukyu and Mainland populations were placed in separate groups, also revealed that although all the three levels of hierarchy were significant (all *P* < 0.001), the majority of variation was attributable to among groups (*F*_CT_ = 0.871; Table S6), indicating that there is a strong genetic differentiation between the native and invasive populations.

The STRUCTURE analysis revealed the most likely number of genetic clusters over all the sampled area to be *K* = 2 (Table S7). The two genetic clusters exactly corresponded to Ryukyu and Mainland populations, with all individuals assigned to the expected cluster with almost 100% probability (Fig. 3). The NJ tree detected the same trend, in which two discrete clades were exhibited with a long distance between the clades of Ryukyu and Mainland populations (Fig. 4). Although the mitochondrial haplotypes also showed two separate clades (Fig. 2), these were independent from the phylogeny constructed with microsatellite data. In the individual-level NJ tree, the clades of the mitochondrial haplotype did not form monophyletic groups or show any signs of divergence (Fig. 4b). This suggests that genetic variation in mitochondrial and nuclear microsatellite DNA do not correlate with each other.

**Fig. 3.**
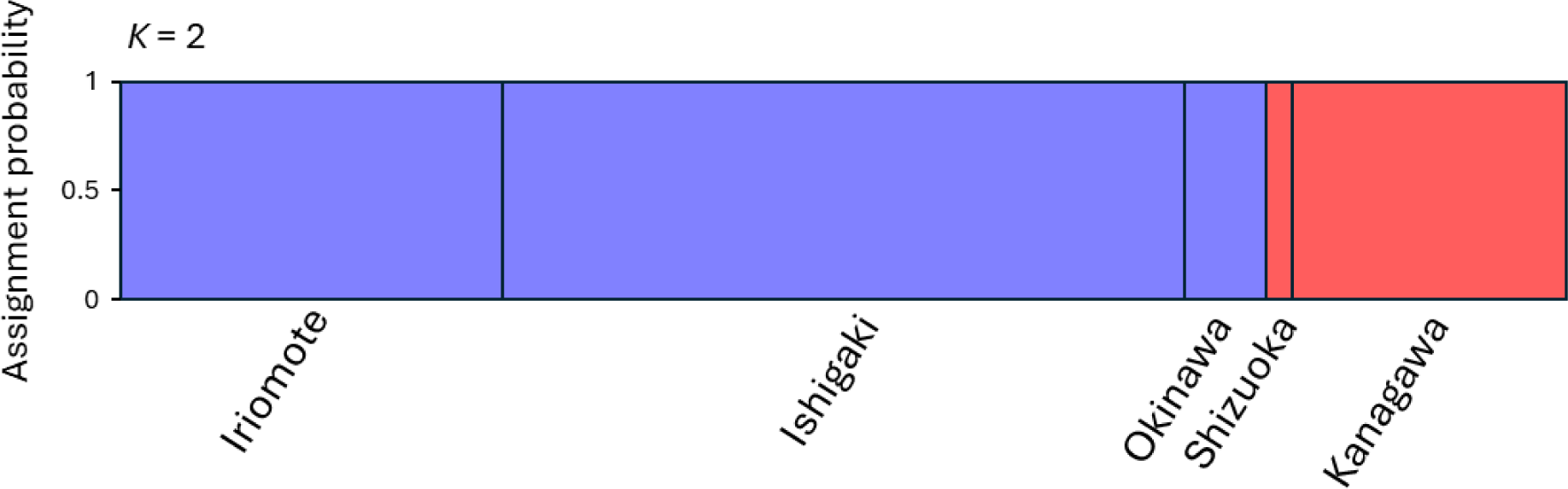
Population genetic structure of five populations in *Melittobia sosui* estimated by a Bayesian clustering analysis with STRUCTURE based on 53 polymorphic microsatellite DNA markers. Inferred assignment probability that individuals belong to each genetic cluster is represented, when the number of clusters is the most likely value based on the method of D*K* (*K* = 2).

**Fig. 4.**
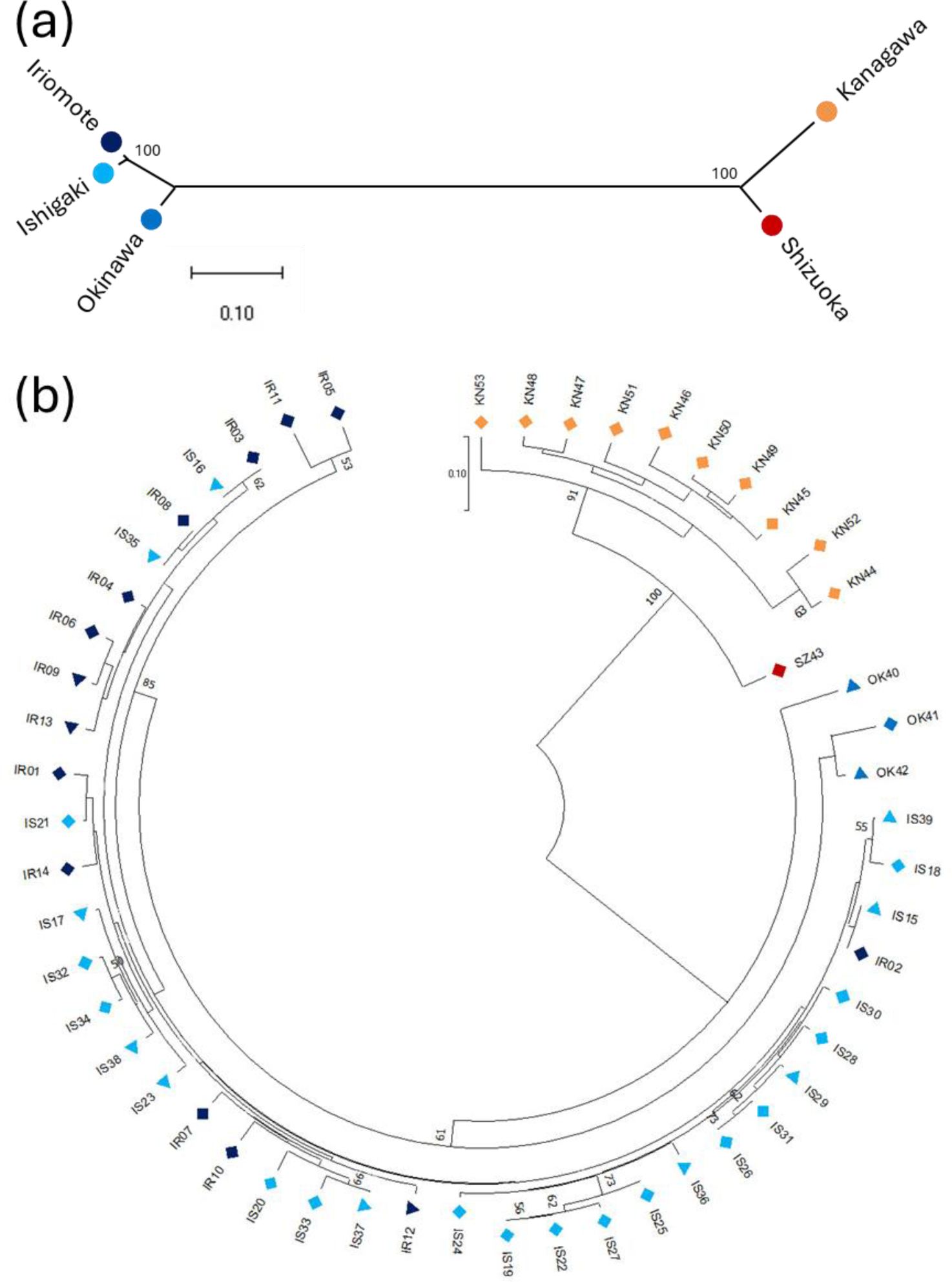
A population-level neighbor-joining tree (a) and an individual-level neighbor-joining tree (b) of *M. sosui* from the native (Iriomote, Ishigaki, and Okinawa) and the invasive (Shizuoka and Kanagawa) populations, based on 53 polymorphic microsatellite DNA markers. The colors of the symbols indicate the population sampled: dark blue, Iriomote (IR); light blue, Ishigaki (IS); blue, Okinawa (OK); red, Shizuoka (SZ); orange, Kanagawa (KN). The shape of the symbols in b indicates mitochondrial haplotype clades: diamond, left clade; triangle, right clade in Fig. 2. Numbers at each node indicate bootstrap values (only ≥ 50% values are shown). The scales bar indicates genetic distance.

The genetic diversity indices calculated based on microsatellite loci were in the same range and not significantly different between Ryukyu and Mainland populations (Table 2 and S8; Wilcoxon signed rank test: number of alleles, *P* = 0.21; allelic richness, *P* = 0.89; gene diversity, *P* = 0.70). On the average over the 53 microsatellite loci examined, only 19.6% of alleles were shared with both population groups, while remaining 42.5% and 37.8% were private alleles that only present in the Ryukyu and Mainland population groups, respectively (Table S9). Even in Mainland populations, which were regarded as new habitat area, relatively high proportion of private alleles were found, and there was no significant difference in the frequencies between the population groups (Wilcoxon signed rank test, *P* = 0.21).

**Table 2.**
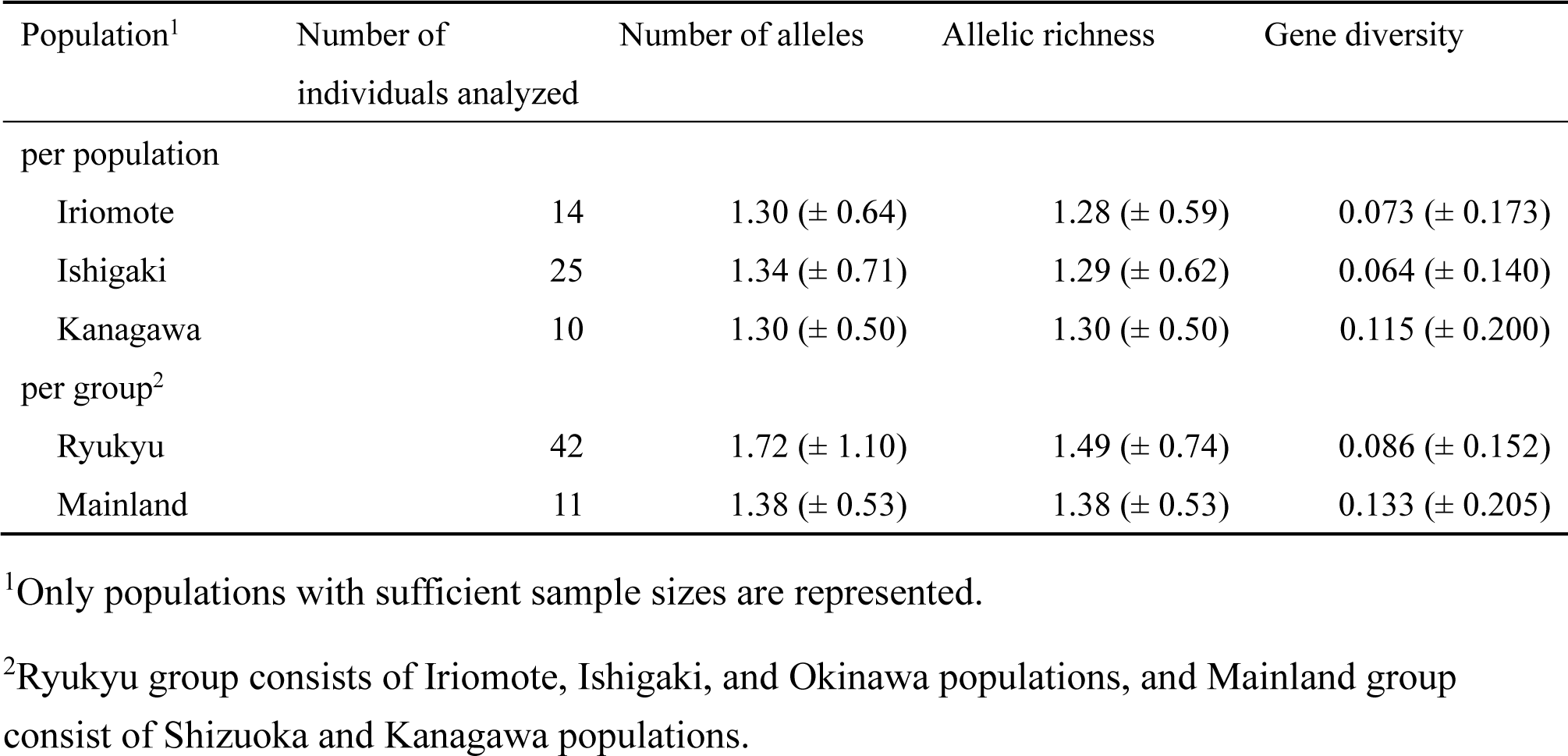
Genetic diversity indexes (mean ± SD) of 53 polymorphic microsatellite DNA markers in *Melittobia sosui*.

### PCR detection for Wolbachia infection

An amplified product of expected size was not detected from all the samples examined, in contrast to the positive control of *M. australica*, suggesting that the sampled populations of *M. sosui* in this study are not infected by *Wolbachia*.

## Discussion

We examined the population genetic structure and genetic diversity within and between native and invasive areas in *M. sosui*, using mitochondrial and nuclear microsatellite DNA markers. Generally, invasive populations are expected to show reduced genetic diversity due to bottlenecks associated with founder events. Present study showed that there was no variation in mitochondrial DNA in the invasive Mainland populations, with only one haplotype which was nested within variation in the haplotypes found in the native Ryukyu populations. However, there were considerable variations in microsatellite DNA in Mainland populations, which were comparable with ones in Ryukyu populations. Moreover, in Mainland, 45.5% microsatellite loci had private alleles, which were not found in Ryukyu populations. These results suggest that Mainland populations could have diverged from Ryukyu populations, but the timing of the divergence was not recently.

One of possible explanations for the high genetic diversity in microsatellite DNA in the invasive populations could be introduction by multiple individuals. At the mitochondrial DNA level, all the individuals shared only one haplotype in Mainland populations, but we cannot distinguish whether the colonization originated from only a single female, multiple females with the same haplotype, or multiple females with different haplotypes but one became dominant due to genetic drift or selection. However, irrespective of the number of females for the origins of Mainland populations, the high genetic diversity in microsatellite DNA in the mainland was caused by the high frequency of private alleles. The high divergence was likely to arise as novel mutations after it diverged from Ryukyu populations.

The contrast between no mitochondrial variation and substantial microsatellite variation found in Mainland populations may be explained by the difference in mutation rate between mitochondrial and microsatellite DNA. In general, mutation rate is higher in microsatellite than in mitochondria. Mutation rate in mitochondrial COI region was estimated to be 7.4 × 10^−5^ per generation in the parasitoid wasp *Nasonia* (Raychoudhury et al. 2010), which belongs to the same superfamily with *Melittobia*. Microsatellite mutation rate is typically in the order of 10^−3^ to 10^−5^ per locus per generation (Ellegren 2004; Chapuis et al. 2015), and falls within the range in other Hymenopteran species (Moritz et al. 2003; Schmid-Hempel et al. 2007). Although mutation rate may vary depending on the effective population size and the population structure of the species, the proportion of microsatellite loci at which mutation did not occur during generations (*t*) can simply be calculated with mutation rate per locus per generation (*m*) as follows:

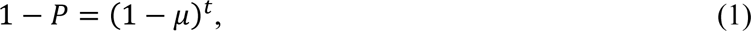

where *P* is the proportion of loci with mutated alleles. Given *m* = 1.0 × 10^−3^ to 1.0 × 10^−5^ and *P* = 0.455, the divergence time between Mainland and Ryukyu populations is estimated to be 6.1 × 10^2^ to 6.1 × 10^4^ generations ago. This implies that mutations could arise at the observed proportion of microsatellite loci, before one mutation occurs in mitochondrial COI region. In the meanwhile, although genetic diversity is generally expected to be smaller in invasive populations than in native populations, Mainland populations had a similar level of genetic diversity in microsatellite DNA with Ryukyu populations. This could be explained because the mutation rate of microsatellite DNA is so high that genetic diversity will converge at the same level for a long time (Nauta and Weising 1996).

Along with the generally high mutation rate in microsatellite DNA, the parasitic life history of *Melittobia* species may facilitate the accumulation of newly emerged mutations. Parasitic lifestyle was expected to elevate nucleotide substitution rate due to small effective population sizes associated with frequent founder events (Page et al. 1998; Kaltenpoth et al. 2012). In the case of *Melittobia*, in which individuals mate within their natal broods, and closely related females often lay eggs on the same host patches, inbreeding will continue to occur unless more than one unrelated females disperse to the same host to lay eggs (Matthews et al. 2009; Abe and Pannebakker 2107; Abe et al. 2021). Actually, *M. sosui* experienced strong inbreeding as shown in the present study. Populations with smaller effective population size will experience faster rate of substitution by genetic drift (Ohta 1992). If mutation occur at a small inbreeding lineage, the mutation will be fixed with a higher probability than in a large non-structured population. Different mutations are likely to be fixed at each inbreeding lineage, resulting in a greater genetic diversity in the whole population. Analogously, population subdivision into genetically isolated lineages and subsequent admixture were expected to lead to an increased genetic divergence (Frankham et al. 2002; Hartl and Clark 2007). Although this kind of effect will have influences on both mitochondrial and microsatellite DNA, it is likely to accelerate the accumulation of once-emerged mutations in the populations.

Another possibility for the reduced variation in mitochondrial but not in nuclear microsatellite DNA could be a cytoplasmic sweep by maternally coinherited *Wolbachia*. A similar pattern to the present study has been reported from the parasitoid wasp *Nasonia vitripennis*, in which North American populations had much smaller microsatellite variation than Europe populations, but had a similar level of microsatellite variation (Raychoudhury et al. 2010). However, in contrast to the present study, the microsatellite data showed shared polymorphisms and did not split between the two population groups. This excludes the possibility that the microsatellite diversity has been newly acquired after the divergence, instead indicates that the mitochondrial variation is likely to be decreased by *Wolbachia*-induced sweep. Although the present study showed that *M. sosui* was not infected by *Wolbachia*, other maternally inherited bacteria or genetic elements could be the cause of the observed reduced variation in mitochondrial DNA. However, so far, there is no evidence suggesting that the reproduction of *M. sosui* is manipulated by any genetic elements. All the broods collected in the present study and laboratory strains in *M. sosui* always had at least one male, ruling out parthenogenesis. *Melittobia* species originally show female-biased sex ratio for their own adaptive reproduction (Abe et al. 2003; Matthews et al. 2009; Abe et al. 2021), suggesting that male-killing and feminization are not likely to yield additional benefits. Hybridization between a Mainland (Kanagawa) strain and a Ryukyu (Ishigaki) strain, which were derived from the different mitochondrial clusters (Fig. 2), produced viable offspring in both male and female directions, implying that cytoplasmic incompatibility is not caused at least in the combination of the strains. Eventually, with the current data, we cannot eliminate the possibility of sweep by an adapted mitochondrial haplotype itself, for which it would be beneficial to compare the fitness consequences of each strain for future studies.

Based on the observational data in the previous literature (van den Assem and Maeta 1980; van den Assem et al. 1982; Maeta 1985; Maeta et al. 1996) and the present data from the field, we expected that the population of *M. sosui* was founded in the Japanese mainland by an individual(s) migrated from Ryukyu area after 1980s. However, although the present mitochondrial data supported this hypothesis, the microsatellite data contradicted this. According to equation (1) with the upper limit of microsatellite mutation rate (*m* = 1.0 × 10^−3^) and that *Melittobia* species have about 6 generations per year in Japanese mainland (Maeta 1978), Mainland populations of *M. sosui* was estimated to diverge from Ryukyu populations at least about 100 years ago.

Then, how did *M. sosui* invade and settle in the mainland? One possibility is that *M. sosui* had already invaded in the mainland over 100 years ago and was distributed at low densities. If the density was low, it might not be collected in the previous studies. Even in small populations, *Melittobia* species can sustain the population through repeating inbreeding. Furthermore, as discussed above, if small populations were distributed at low densities and subsequently mixed, it may have promoted an increase in genetic diversity throughout the population. For alternative possibility, there was a ghost population, which diverged from Ryukyu populations over 100 years ago. This population might have formed in regions where previous study on *Melittobia* were not conducted vigorously. The ghost population might have gradually expanded its distribution, and reached Shizuoka and Kanagawa in the mainland without experiencing bottleneck events. For future study, collecting samples from regions not investigated in the present study, as well as comparing the physiological characteristics, specifically in response to temperature, between Ryukyu and Mainland populations would be useful to examine above possibilities.

In conclusion, the present study found reduced mitochondrial diversity and high microsatellite diversity in the invasive Japanese mainland populations of *M. sosui*. The reduced mitochondrial diversity suggested that the mainland populations were founded by one or a part of individual(s) migrated from the native area. Although the high microsatellite diversity might be explained by introduction by multiple individuals, the observed high diversity resulted from the high proportion of private alleles, which instead suggested that the diversity has been acquired after the population diverged from the native population. Although the present study, which had insufficient sampling areas, could not fully identify the detailed introduction history of *M. sosui* to the Japanese mainland, it can conclude that the mainland populations did not recently diverge from the native populations. More generally, the present results emphasize the usefulness of using molecular techniques, which could lead to different conclusions contrast to observational or historical data.

## Acknowledgements

We thank Mai Miyata, Yoshitaka Kamimura, and Momoko Ichinokawa for valuable discussion for the study. This research was supported by JSPS KAKENHI (grant no. 21K06353 and 21KK0267) to J. A.

